# Lis1 activates dynein motility by pairing it with dynactin

**DOI:** 10.1101/685826

**Authors:** Mohamed M. Elshenawy, Emre Kusakci, Sara Volz, Janina Baumbach, Simon L. Bullock, Ahmet Yildiz

## Abstract

Lissencephaly-1 (Lis1) is a key cofactor for dynein-mediated intracellular transport towards the minus-ends of microtubules (MTs). It remains unclear whether Lis1 serves as an inhibitor or an activator of mammalian dynein motility. Here we use single-molecule imaging and optical trapping to show that Lis1 does not directly alter the stepping and force production of individual dynein motors assembled with dynactin and a cargo adaptor. Instead, Lis1 binding releases dynein from its auto-inhibited state and thereby promotes the formation of an active complex with dynactin. Lis1 also favors recruitment of two dyneins to dynactin, resulting in increased velocity, higher force production and more effective competition against kinesin in a tug-of-war. Lis1 dissociates from motile complexes, indicating that its primary role is to orchestrate the assembly of the transport machinery. These results provide a mechanistic explanation for why Lis1 is required for efficient transport of many dynein-associated cargoes in cells.

---

Cytoplasmic dynein (dynein hereafter) is an AAA+ motor responsible for nearly all motility and force generation towards the microtubule minus-end^1–3^. Dynein is involved in a wide variety of cellular functions, such as positioning of intracellular organelles, breakdown of the nuclear envelope and assembly of the mitotic spindle during mitosis^4–6^. The partial loss of dynein function has been implicated in a range of neurogenerative and neurodevelopmental conditions, including spinal muscular atrophy, amyotrophic lateral sclerosis, Alzheimer’s disease and schizophrenia^7–9^.

The dynein complex (1.4 MDa) is a homodimer of two heavy chains, which recruit smaller associated polypeptides^10^. The C-terminal motor domain of the heavy chain is a catalytic ring of six AAA modules (AAA1-6). ATPase activity at AAA1 powers dynein motility^11, 12^, while AAA3 regulates MT attachment and force generation cycles^13–15^. In contrast to kinesin, whose MT interface is located on the surface of the ATPase core, dynein’s MTBD is separated from the catalytic domain by a coiled-coil stalk^16^. Nucleotide-dependent conformational changes of the linker drive the motility towards the MT minus-end^17, 18^. The tail domain dimerizes the heavy chains^19–21^, and binds a light intermediate chain (LIC) and an intermediate chain (IC) complexed with three light chains (LCs: LC7, LC8 and Tctex)^1, 22, 23^.

When dynein is not bound to its cargo, it forms two distinct conformations, the phi-particle and open conformation, both of which move poorly along MTs^24, 25^. In the phi conformation, two motor domains self-dimerize through interactions between their linker, AAA+ ring and stalk regions and weakly interact with MTs. In the open conformation, the motor domains are more flexible and point towards each other, which is unfavorable for processive motility^24, 26^. Dynein interacts with its activating cofactor dynactin and binds to its cellular cargos in the open conformation. Transitions between the phi and open conformation are proposed to be an important part of dynein regulation^24, 26^, but the molecular cues that govern this transition remain unclear.

Dynein and dynactin are recruited to cargos through coiled-coil adaptor proteins in a mutually dependent manner (Fig. 1a)^27–29^. Formation of dynein-dynactin-cargo adaptor complex (DDX) aligns the dynein motor domains in a parallel conformation and activates processive motility along MTs^30, 31^. These adaptors recruit dynein to a specific set of cargos^29, 32^, enabling a single dynein gene to be responsible for nearly all minus-end directed functions along MTs. Members of the BicD family, BicD2 and BicDR1, are well-characterized coiled-coil adaptors that link dynein to Golgi-derived Rab6 vesicles, as well as nuclear pore complexes and viruses^31, 33, 34^. Structural studies showed that BicDR1 recruits two dyneins to dynactin, while the N-terminal coiled-coil domain of BicD2 (BiCD2N) mostly recruits a single dynein^35, 36^. The increased probability of recruiting two dyneins per dynactin results in complexes assembled with BicDR1 (DDR) moving faster and producing more force than complexes formed with BicD2N (DDB)^35^. The differences elicited by BicD2 and BicDR1 in dynein motility may play a critical role in sorting of Rab6 vesicles during neuronal differentiation^28, 33^.

**Figure 1.**
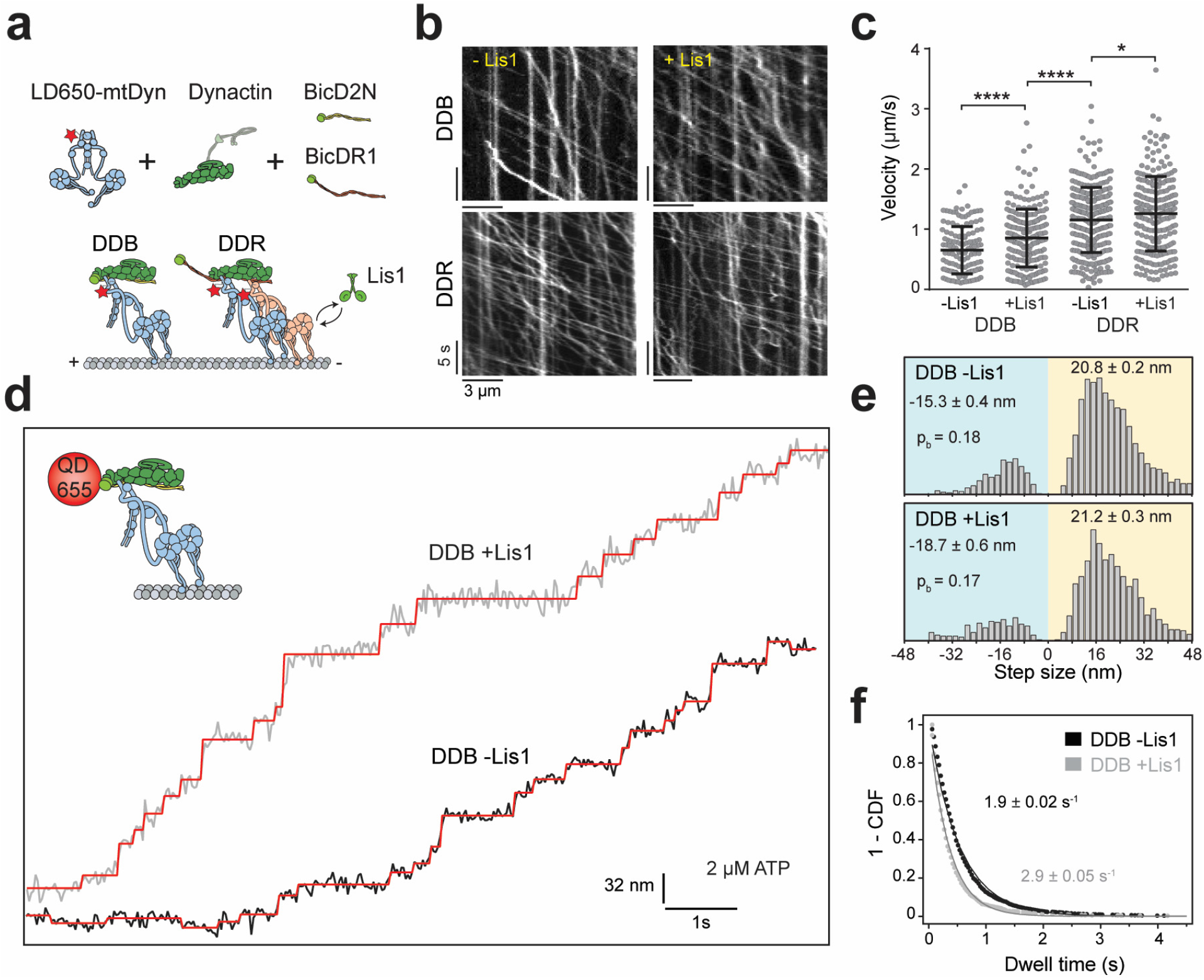
Lis1 increases the stepping rate of dynein-dynactin. **(a)** Schematic depiction of DDB and DDR complexes and Lis1. All three components (dynein, dynactin and a cargo adaptor) must be present for processive motility of the complex. BicD2N primarily recruits a single dynein to dynactin, whereas BicDR1 recruits two dynein. Lis1 binds to the dynein motor domain. **(b)** Sample kymographs show the motility of DDB and DDR on MTs. **(c)** Velocity distribution of DDB and DDR with and without 600 nM Lis1. The center line and whiskers represent the mean and s.d., respectively (*n* = 209, 213, 307, and 241 from left to right, ****p < 0.0001, *p <0.05, two-tailed t-test). (**d**) Representative stepping traces of DDB at 2 μM ATP along the longitudinal axis of the MT. (Insert) Dynein was labeled with a QD (red circle) and its motility was tracked at 30 ms temporal resolution. Red staircases represent a fit of the traces to a step finding algorithm. (**e**) Normalized histograms of DDB step size (*n* = 2,076 steps for DDB -Lis1 and 1,374 for DDB +Lis1). Average forward and backward step sizes and the probability of backward stepping (*p*_*b*_) are shown (± s.e.m.). (**f**) Inverse cumulative distribution of dwell times between consecutive steps along the longitudinal axis. Solid curves represent fitting to an exponential decay (decay rate ± s.e., *n* = 2138 for DDB -Lis1 and 1441 for DDB +Lis1).

Dynein motility is also regulated by Lis1, which directly interacts with the dynein motor domain^37^. Lis1 inhibition reduces transport of a wide variety of cargoes in eukaryotic cells, including endosomes, lysosomes, mRNAs, centrosomes and nuclei^38–45^. The critical role of Lis1 is underscored by the discovery that haploinsufficiency of Lis1 gene causes a smooth brain disorder (lissencephaly) in humans, which is associated with a failure of nuclear migration^46^. Lis1 also forms a homodimer, with each monomer comprising of an N-terminal dimerization domain and a C-terminal β-propeller domain that binds dynein at the interface between AAA3/4 (Fig. 1a)^37, 47^.

The mechanism by which Lis1 regulates dynein motility is controversial. *In vitro* studies on yeast dynein revealed that a Lis1 homolog, Pac1 increases MT affinity, blocks nucleotide-dependent remodeling of the linker domain, and significantly reduces dynein velocity^37, 48^. However, the view of Lis1 as an inhibitor of dynein is counterintuitive since Lis1 promotes dynein-mediated cargo transport in many cell types and organisms^38–45, 49^. Studies on isolated mammalian dynein proposed that Lis1 transiently interacts with dynein, enhances dynein’s affinity to MTs on high-load cargos by inducing a persistent-force generation state^41, 50^. However, Lis1’s function is not restricted to high-load cargos, and it is also required for the transport of smaller cargos^40, 42, 43, 45^. These studies were performed before it was understood that isolated dynein motors are autoinhibited in the absence of dynactin and a cargo adaptor, and may not reflect the force generation mechanism of active DDX complexes^29, 32, 51^. *In vivo* studies gave rise to models that Lis1 is only required for targeting dynein to the MTs, with dissociation of Lis1 triggering the initiation of transport^38, 42, 52–54^ and that Lis1 promotes the interaction of dynein and dynactin^43, 44^. Consistent with these models, recent *in vitro* studies showed that mammalian Lis1 can increase the frequency and velocity of DDB motility^54–56^, but the underlying mechanism remained unknown.

In this study, we determined the effect of Lis1 binding on the motility, stepping, and force generation of DDB and DDR using single-molecule imaging and optical trapping *in vitro*. We found that Lis1 has no significant effect on the force generation of single dyneins after they have associated with dynactin and the cargo adaptor. Instead, Lis1 binding promotes the assembly of dynein to dynactin by releasing dynein from its phi conformation to open conformation. Lis1 also favors the association of two dyneins to dynactin, and this accounts for the increase in both velocity and force generation. As a result, the presence of Lis1 shifts the force balance towards dynein’s direction during a tug-of-war with a plus-end directed kinesin. Our work reveals how binding of Lis1 activates the motility of mammalian dynein and is thereby required for efficient transport of cargos in cells.

## Results

### Lis1 increases the stepping rate of DDX complexes

We first tested the effect of human Lis1 binding on the velocity of DDB and DDR complexes assembled with wild-type human dynein (wtDyn). In agreement with previous measurements^55, 56^, DDB moved 30% faster in 600 nM Lis1 (two-tailed t-test, *p* = 10^−4^). We also observed a modest (10%) increase in DDR velocity by Lis1 addition (Supplementary Fig. 1). Similar results were obtained using a dynein mutant (mtDyn) that disfavors the phi conformation^26^ (Fig. 1a-c, and Supplementary Movies 1 and 2). Because mtDyn increases the number of active DDX complexes^26^, we used this construct hereafter to study the effect of Lis1 on dynein stepping and force generation. To distinguish whether Lis1 binding increases dynein step size or stepping rate for faster movement, we determined the stepping behavior of DDB in the presence and absence of Lis1 at limiting (2 μM) ATP concentrations. We labeled dynein with a bright quantum dot (QD) at its N-terminus and tracked the motility of single DDB complexes at nanometer precision. In the absence of Lis1, DDB has highly variable step size, frequently taking backward steps (Fig. 1d). We found that Lis1 addition did not alter the size and direction of steps taken by dynein. Instead, DDB stepped more frequently in Lis1 (2.9 ± 0.05 vs. 1.9 ± 0.02s^−1^, ± s.e.; two-tailed t-test, *p* = 10^−13^; Fig 1d-f and Supplementary Figure 2). We concluded that Lis1 increases the stepping rate, not the mean step size, which accounts for the faster velocity of dynein.

### Lis1 increases the force production of DDX complexes

We tested whether Lis1 affects the force generation of dynein when this motor forms an active complex with dynactin and a cargo adaptor using an optical trap. We sparsely coated polystyrene beads with BicD2N and BicDR1 adaptors and assembled DDB and DDR complexes on beads. This assay geometry ensures that beads are driven by active dynein motors assembled to cargo adaptors and eliminates the possibility of cargo adaptor multimerization. Using a fixed trap, we observed that the stall force of DDB increases by 22% in 600 nM Lis1 (4.1 ± 0.1 vs. 5.4 ± 0.1 pN, mean ± s.e.m; two-tailed t-test, *p* = 10^−11^; Fig. 2a-c). Lis1 also resulted in an increase in DDR force production, although the magnitude of the effect was more modest (5.4 ± 0.1 vs. 6.1 ± 0.1 pN; two-tailed t-test, *p* = 10^−4^; Fig. 2c). Contrary to a previous study on isolated dyneins^50^, we did not observe a significant difference in the stall duration of DDB and DDR in the presence of Lis1 (Fig. 2d and Supplementary Fig. 3).

**Figure 2.**
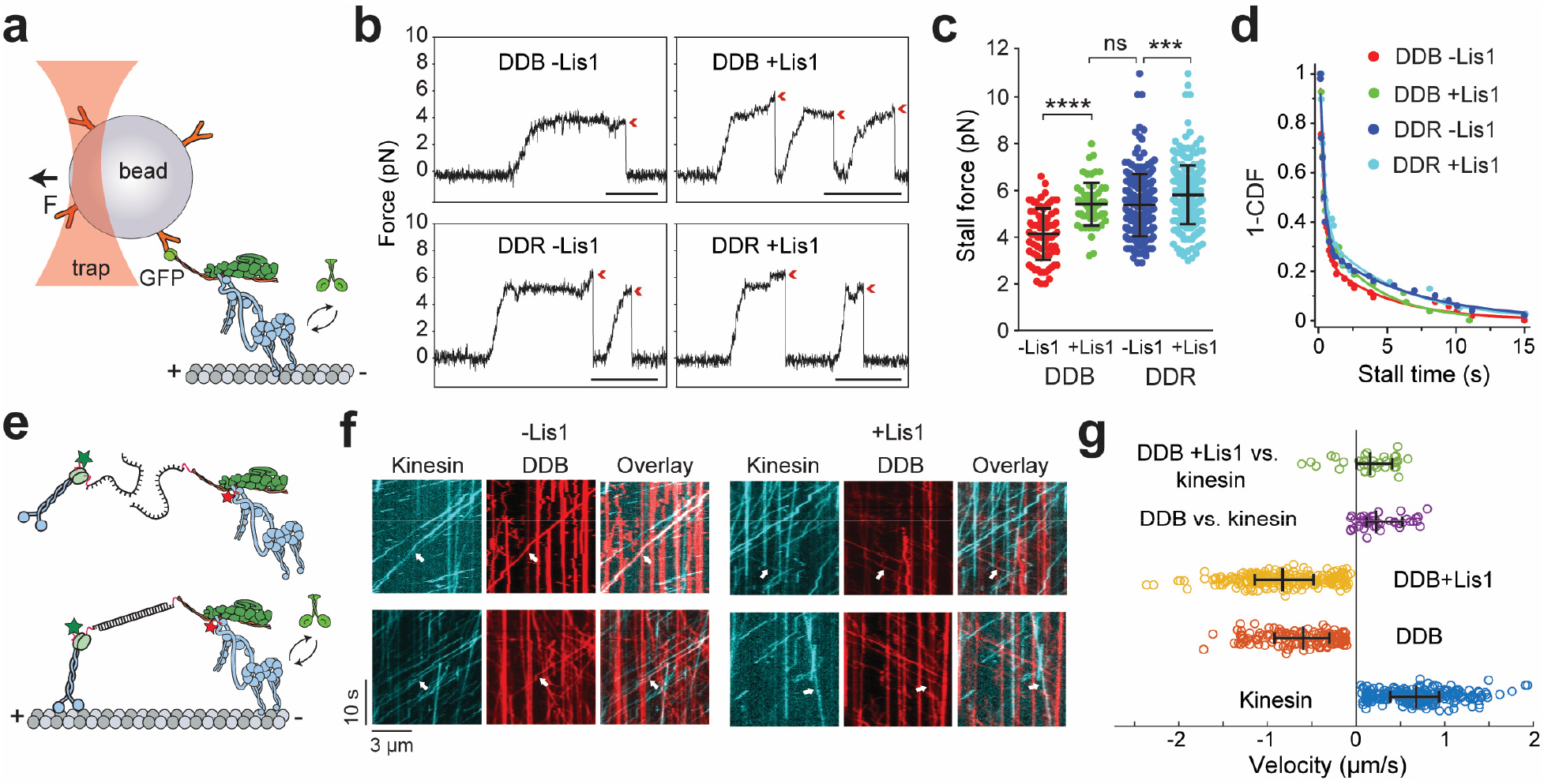
Lis1 increases the force production of dynein-dynactin. **(a)** Schematic of a fixed optical trapping assay for measuring the dynein stall force. **(b)** Typical stalls of beads driven by a single DDB or DDR. Red arrowheads denote the detachment of the motor from the MT after the stall event. Scale bars are 1 s. **(c)** Distribution of motor stall forces in the absence and presence of 600 nM Lis1. The center line and whiskers represent the mean and s.d., respectively (*n* = 80 from 19 beads, 61 from 15 beads, 212 from 38 beads, and 152 from 32 beads from left to right, ns: non-significant, ****p < 0.0001, ***p <0.005, two-tailed t-test). **(d)** Inverse cumulative distribution of stall durations of DDB and DDR in the presence and absence of Lis1. Solid curves represent fitting to a two-exponential decay (decay time ± s.e., n = 53 for DDB -Lis1, 27 for DDB +Lis1, 50 for DDR -Lis1, and 39 for DDR +Lis1). **(e)** Schematic depiction of the *in vitro* tug-of-war assay. DDB and kinesin were labeled with different-colored fluorescent dyes and tethered using a DNA scaffold. **(f)** Representative kymographs show the motility of LD650-dynein (red) and TMR-kinesin (cyan). White arrows show DDB-kinesin colocalizers. **(g)** Velocity distribution of DDB, kinesin, and DDB-kinesin assemblies in the absence and presence of Lis1. The center line and whiskers represent the median and 65% CI, respectively (*n* = 33, 45, 193, 179, and 210 from top to bottom). Negative velocities represent movement towards the MT minus-end.

We then tested whether the increase in DDB force production by Lis1 also increases the likelihood of DDB to win a tug-of-war against a plus-end-directed kinesin-1. We labeled dynein and kinesin with different fluorescent dyes and pitted one DDB against one kinesin using a DNA tether. Consistent with our previous measurements^51^, the majority (87%) of kinesin-DDB assemblies move towards the plus-end at a median velocity of 185 nm/s in the absence of Lis1. Addition of Lis1 slowed down the mean velocity of plus-end-directed movement by 40% and increased the percentage of complexes moving towards the minus-end from 13% to 22% (Fig 2e-g and Supplementary Movie 3). Collectively, these results demonstrate that Lis1 increases the force production of DDX complexes and shifts the force balance more in dynein’s favor in a tug-of-war.

### Lis1 has no effect on force generation of single dynein when bound to dynactin and a cargo adaptor

We next turned our attention to understanding how Lis1 increases the velocity and force production of DDB and DDR. To test whether Lis1 binding alters mechanochemical properties of single dynein assembled with dynactin, we mixed LD650-labeled full-length dynein (mtDyn) and a TMR-labeled dynein tail construct (Dyn_LT_, containing residues 1–1,074 of the heavy chain and associated chains). Because Dyn_LT_ lacks the motor region, processive runs of this construct can only be achieved through its side-by-side recruitment with mtDyn to dynactin. We measured the velocity of dual-labeled complexes that contain both mtDyn and Dyn_LT_ in the presence of BicD2N (DTB) and BicDR1 (DTR). We found that Lis1 addition has no effect on the mean velocity of complexes containing a single dynein (Fig. 3a-c and Supplementary Movie 4).

**Figure 3.**
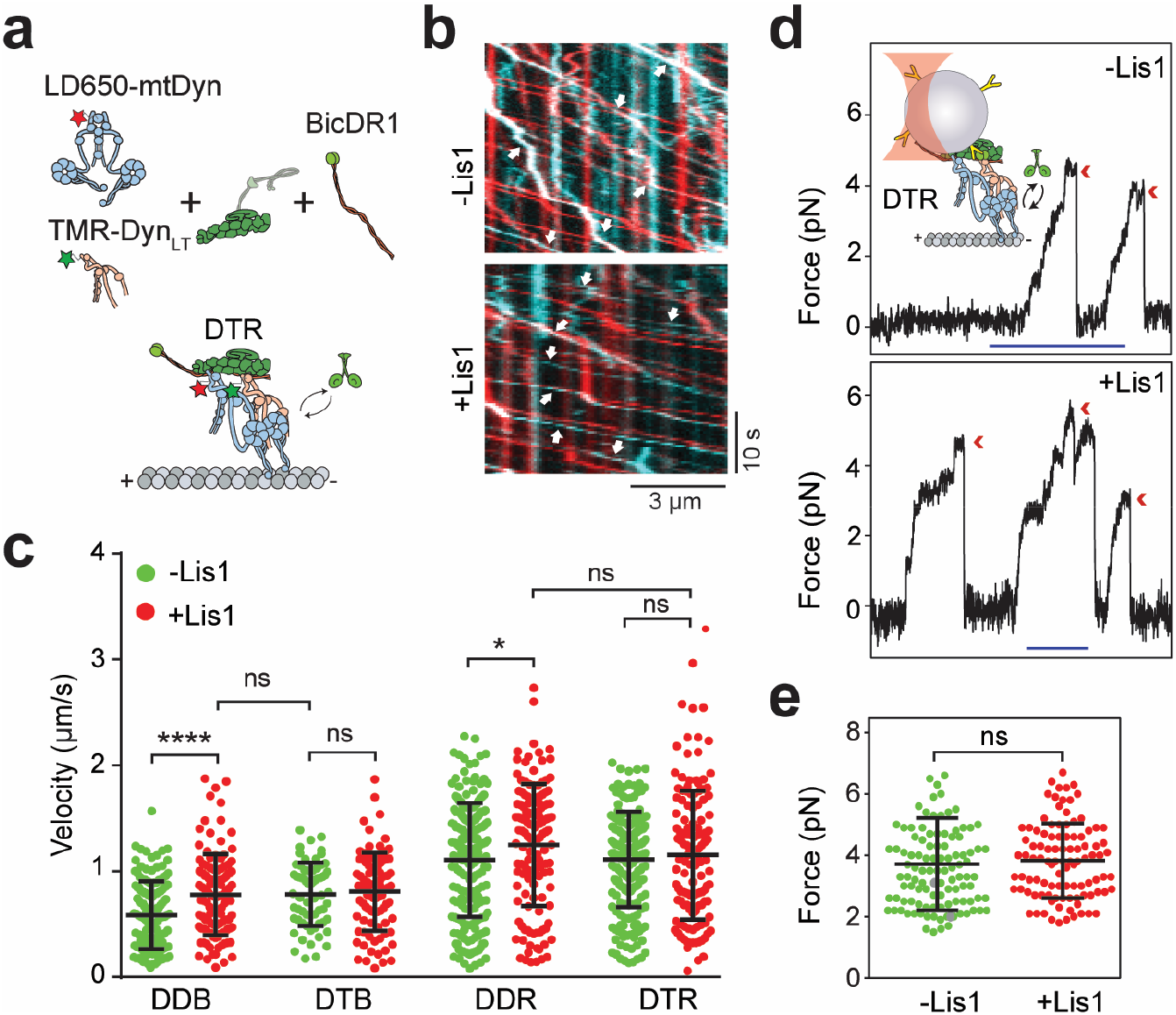
Lis1 does not affect the force generation and velocity of a single dynein complexed to dynactin and a cargo adaptor. **(a)** Schematic depiction of the DTR complex. Full-length dynein and Dyn_LT_ are labeled with LD650 and TMR dyes, respectively. **(b)** Representative kymographs show the motility of Dyn (red) and Dyn_LT_ (cyan). White arrows represent colocalization of TMR and LD650. **(c)** Velocity distribution of DDB, DTB, DDR and DTR in the presence and absence of Lis1. The center line and whiskers represent the mean and s.d., respectively (*n* = 144, 117, 65, 88, 209, 134, 213, and 126 from left to right, ns: non-significant, ****p < 0.0001, *p < 0.05, two-tailed t-test). (**d**) Typical stalls of beads driven by DTR in the absence and presence of Lis1. Red arrowheads denote the detachment of the motor from the MT after the stall event. Scale bar is 1 s. **(e)** Distribution of DTR stall force. The center line and whiskers represent the mean and s.d., respectively (*n* = 111 stalls from 23 beads for -Lis1 and 101 stalls from 21 beads for +Lis1, ns: non-significant, two-tailed t-test).

To test whether Lis1 alters the stepping properties of single dynein bound to dynactin, we tracked beads driven by single DTR under constant 1 pN hindering force exerted by the trap. Unlike DDB, Lis1 addition did not alter the stepping rate of DTR in 1 mM ATP (44.4 ± 0.5 vs. 46.6 ± 0.5 s^−1^, decay rate ± s.e.; two-tailed t-test, *p* = 0.83; Supplementary Fig. 4a-c). Measurements under a fixed trap also revealed that Lis1 addition also does not affect the stall force and stall duration of DTR (Fig. 3d-e and Supplementary Fig. 4d). Therefore, Lis1 does not directly affect the mechanical properties of a single dynein bound to dynactin.

### Lis1 promotes recruitment of two dyneins to dynactin

The effect of Lis1 on DDB velocity and force production is strikingly similar to the recruitment of a second dynein to dynactin^35^, leading us to hypothesize that Lis1 regulates the stoichiometry of dynein per dynactin. This possibility could not be tested directly in our previous experiments, because dynein copy number per complex could not be directly determined from stepping (Fig. 1c) and force (Fig. 2b) measurements. To address this, we mixed TMR- and LD650-labeled dynein with dynactin and BicD2N and quantified the percentage of processive complexes that contained both fluorophores. Lis1 addition increased the percentage of colocalization from 14% to 24% (*p* = 0.018, two-tailed t-test, Fig. 4a-b and Supplementary Movie 5). After correction for labeling efficiency and complexes with two dyneins labelled with the same color, we estimated that Lis1 addition increases the percentage of complexes containing two dyneins from 22% to 42%). Lis1 addition also increased the velocity of single-colored complexes that contain predominantly one dynein motor, but not that of dual-colored complexes that contain two dyneins (Fig. 4c). Similar results were obtained when BicDR1 was used as a cargo adaptor, but the effect of Lis1 addition was more modest, presumably because DDR complexes are already predisposed to contain two dyneins. Collectively, these results demonstrate that Lis1 favors the recruitment of two dynein motors to dynactin, which accounts for the faster velocity of these complexes in the presence of Lis1^35, 55, 56^.

**Figure 4.**
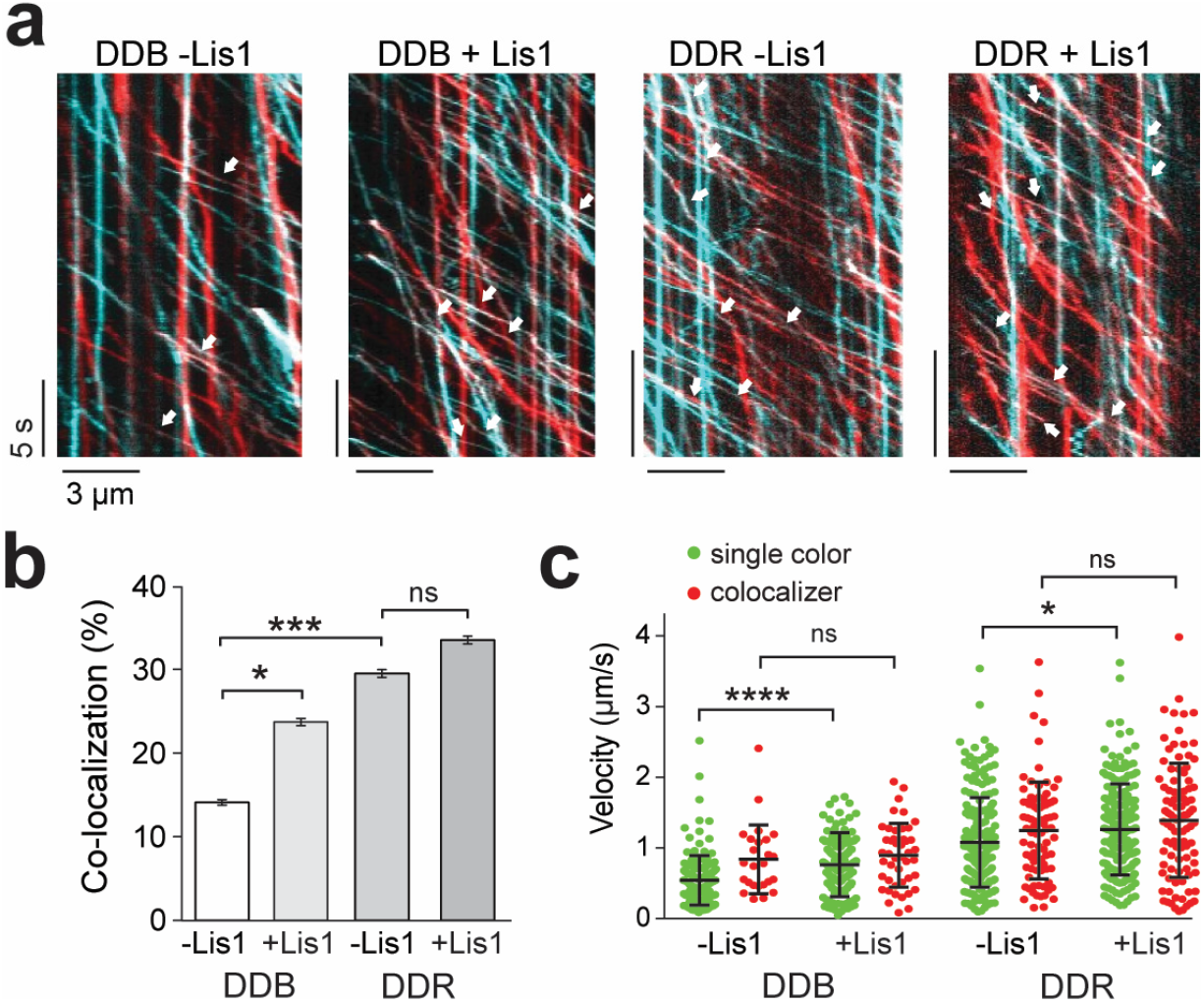
Lis1 favors the recruitment of two dyneins to dynactin. **(a)** Dynein was labelled with TMR and LD650 dyes and mixed with dynactin and cargo adaptors in absence and presence of Lis1. Representative kymographs show the motility of LD650- (red) and TMR- (cyan) labeled dynein. Two colors assemblies are denoted with white arrows. **(b)** The percentage of processive complexes that contain both TMR and LD650 signal (mean ± s.e.m, *n* = 178, 190, 289, 290 from left to right, ns: non-significant, *p < 0.05, ***p <0.005, two-tailed t-test). **(c)** Velocity distribution of single-colored and dual-colored complexes of DDB and DDR in the presence and absence of Lis1. The line and whiskers represent the mean and s.d., respectively (*n* = 153, 25, 145, 45, 204, 85, 193, and 97 from left to right, ns: non-significant, *p < 0.05, ****p < 0.0001, two-tailed t-test).

In addition to binding the motor domain, previous studies reported that mammalian Lis1 interacts with dynein through the tail region of the heavy chain and the intermediate chain, as well as with dynactin at the p50 subunit^57, 58^. To test whether these interactions affect association of dynein with dynactin, we quantified the recruitment of Dyn_LT_ side-by-side with mtDyn to dynactin in the presence and absence of Lis1. Addition of Lis1 had no effect on the ratio of Dyn_LT_ to mtDyn processive runs in both BicD2N and BicDR1 (Supplementary Fig. 5), revealing that Lis1 does not have a direct role in recruitment of dynein tail to dynactin. We concluded that Lis1 favors recruitment of two dyneins to dynactin through its interactions with the dynein motor domain.

### Lis1 dissociates from motile complexes

To determine whether Lis1 remains stably bound when dynein moves along MTs^54–56^, we mixed LD650-mtDyn and TMR-Lis1 in the presence of dynactin and BicD2N, and quantified the velocity of Lis1-bound DDB complexes moving along MTs (Fig. 5a). In these assay conditions, only 10% of the motile DDB complexes contained Lis1, and these complexes had a significantly lower velocity than other DDB complexes (448 ± 38 vs 726 ± 13 nm/s, mean ± s.e.m; two-tailed t-test, *p* = 10^−9^; Fig. 5b-c). In rare occasions, we also observed Lis1, which diffuses on an MT, hops onto a motile DDB complex on the same MT (Supplementary Fig. 6) and slows down the motility^37, 55^. Therefore, Lis1 reduces dynein velocity if it remains bound to DDX complexes after the initiation of processive motility. However, this has a minor effect on the motile properties of the overall dynein population, because Lis1 typically dissociates from motile complexes.

**Figure 5.**
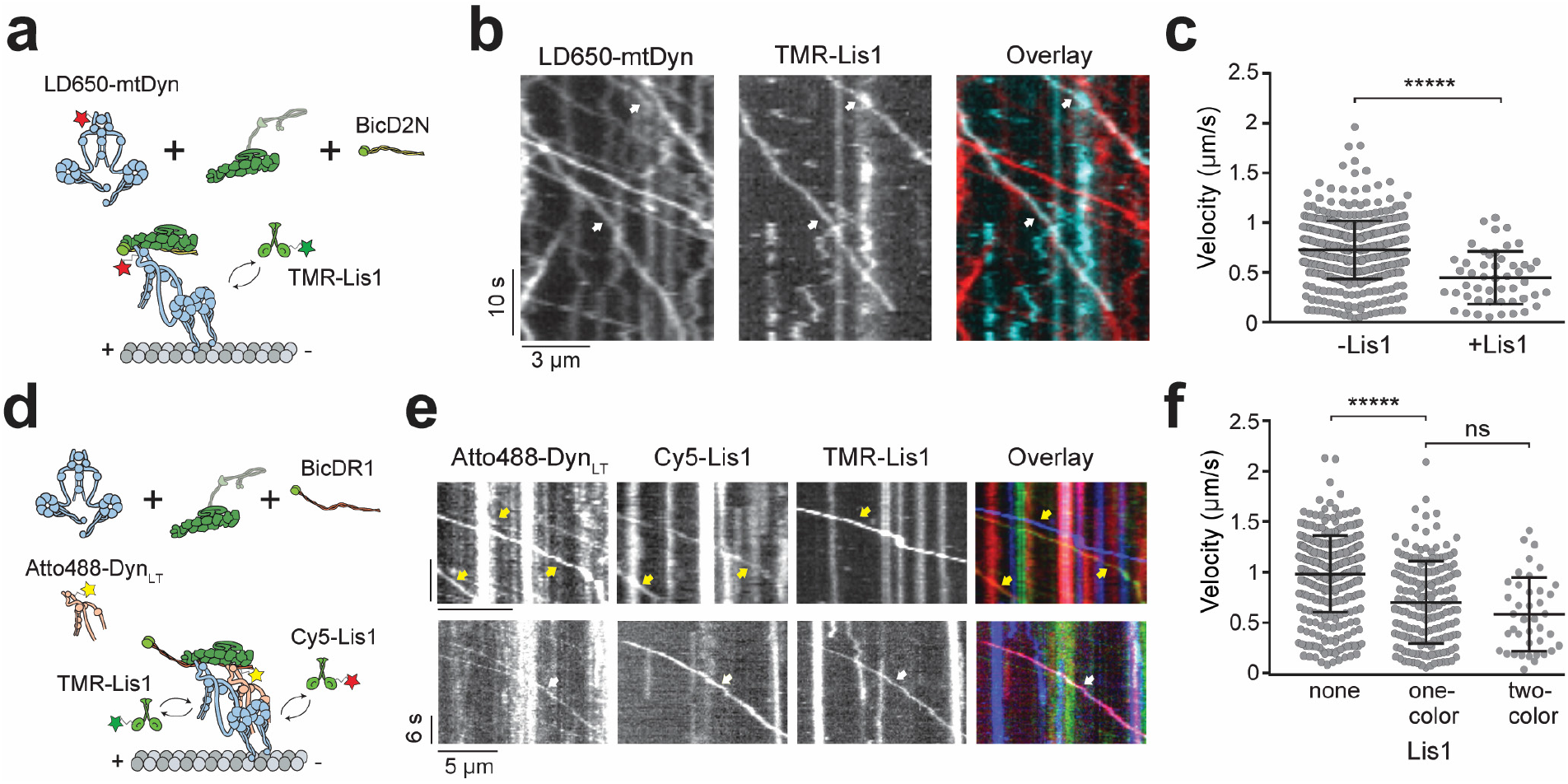
Two Lis1 binds a single dynein and Lis1 binding decreases the velocity of dynein/dynactin. **(a)** Schematic depiction of DDB complex assembled in the presence of 50 nM TMR- Lis1. **(b)** Representative kymographs show the motility of DDB and Lis1 on MTs. White arrows represent colocalization of LD650-Dyn (red) and TMR-Lis1 (cyan). **(c)** Velocity distribution of DDB and DDB- Lis1 assemblies. The center line and whiskers represent the mean and s.d., respectively (*n* = 512, and 49 from left to right, ****p < 0.0001, two-tailed t-test). **(d)** Schematic depiction of DTR complex assembled in the presence of 50 nM TMR- and Cy5-Lis1. **(e)** Representative kymographs show the motility of DTR and Lis1 on MTs. Yellow arrows represent colocalization of Dyn_LT_ (green) and one color of Lis1. White arrows represent colocalization of Dyn_LT_ with Cy5- (red), and TMR-Lis1 (cyan). **(f)** Velocity distribution of DTR that colocalizes with zero, one and two colors of Lis1. The center line and whiskers represent the mean and s.d., respectively (*n* = 357, 172, and 40 from left to right, ns: non-significant, ****p < 0.0001, two-tailed t-test).

Stoichiometry of Lis1 binding to dynein is also not well understood. A previous study reported that DDB can concurrently recruit two Lis1 dimers^55^. However, because Lis1 binding favors recruitment of two dyneins to dynactin, it remains unclear whether each of the two dyneins in DDB binds to Lis1 or a single dynein can simultaneously bind to two Lis1 dimers. To address this, we mixed Atto488-Dyn_LT_ with an equimolar mixture of TMR-Lis1 and Cy5-Lis1 and tested if two Lis1 molecules could bind to the single full-length dynein recruited side-by-side with Atto488-Dyn_LT_. We observed that 7% of complexes that recruit Dyn_LT_ colocalized with both TMR- and Cy5-labeled Lis1 during processive motility and reduced velocity (Fig. 5, Supplementary Fig. 6 and Supplementary Movie 6). Therefore, a single dynein can simultaneously recruit two Lis1 dimers.

### Lis1 promotes the assembly of active DDX complexes

Lis1 stimulates the frequency of minus-end-directed transport under conditions insufficient to induce motility, such as when BicD2N concentration is low^56^. To determine how Lis1 favors initiation of dynein motility when complex formation is strongly limiting, we quantified DDB motility while we lowered the wtDyn concentration 10-, 20-, and 50-fold compared to the standard condition (see Methods). In the absence of Lis1, the percentage of DDB complexes exhibiting motility was decreased at lower dynein concentrations, and motility is almost fully abolished with the 50-fold dilution. Lis1 addition increased the percentage of motility by ~5-fold (Figure 6a-b and Supplementary Movie 7), which is consistent with Lis1 favoring association of dynactin with dynein and the BicD2 orthologue in *Drosophila* cell extracts^43^. However, when we used mtDyn that does not form the phi conformation, we observed robust motility even in the 50-fold dilution condition and no significant increase in the percentage of motile complexes with the addition of Lis1 (Fig. 6a-b). We also mixed equal amounts of TMR- and LD650-labeled dynein with dynactin and BicD2N and quantified the percentage of colocalizing complexes moving along the MTs under limiting dynein conditions. We observed 2% of the processive runs contain both TMR and LD650 signals, indicating that only ~ 5% of complexes are assembled with two dyneins. Additionally, we did not observe an increase in DDB velocity by Lis1 addition under these conditions (Supplementary Fig. 7). This result demonstrates that Lis1 promotes motility by recruiting a single dynein to dynactin under these conditions. Thus, in addition to favoring the association of the second dynein with dynactin, Lis1 also promotes the association of the first dynein to initiate motility.

**Figure 6.**
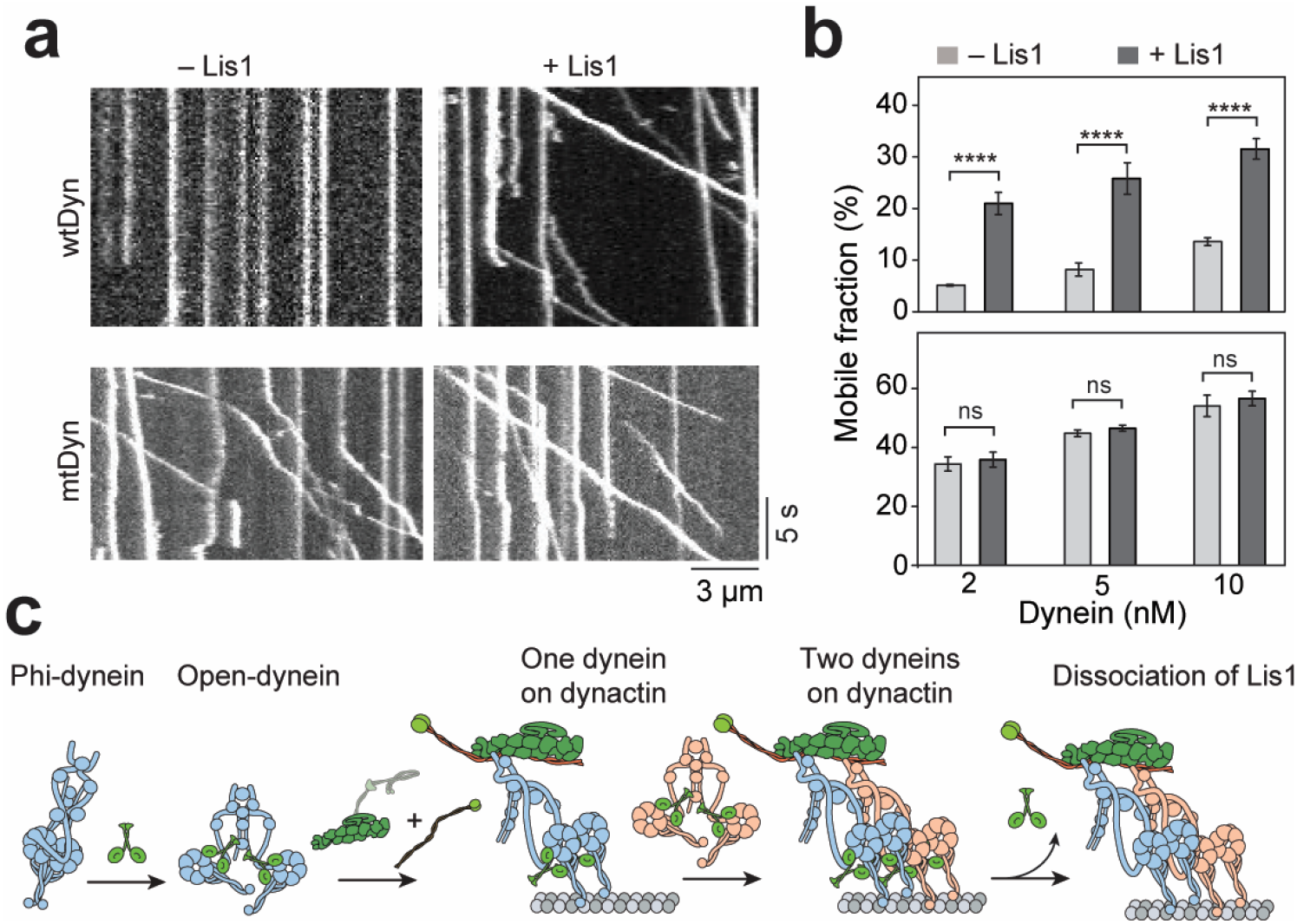
Lis1 promotes assembly of the DDX complex. **(a)** Representative kymographs show the motility of DDB at 5 nM concentration of dynein. **(b)** Ratio comparison of the number of processive runs by DDB to the total number of landed motors on MT (mean ± s.e.m, from left to right *n* = 508, 355, 491, 262, 1244, and 392 for wtDyn, and 234, 459, 426, 1352, 457, and 859 for mtDyn, ns: non-significant, ****p < 0.0001, two-tailed t-test). **(c)** A model for Lis1-mediated assembly of the DDX complex. Lis1 binds to the open-conformation of dynein with one Lis1 dimer for each dynein motor domain. Lis1 binding prevents transitions of the open conformation to the phi conformation, which increases the affinity of dynein to dynactin. Lis1 also facilitates recruitment of a second dynein to the complex resulting in higher force production and faster movement. Lis1 dissociates from these motile complexes after promoting the pairing of dynein with dynactin.

## Discussion

Our results challenge previous views on how Lis1 binding regulates dynein motility. A previous study on isolated dynein suggested that Lis1 binding induces pausing of dynein motility and enhances MT affinity when dynein is subjected to force^50^. Studies on yeast dynein also suggested that Lis1 binding interferes with the powerstroke of the linker domain^48^, suggesting that Lis1 binding reduces dynein force generation. Our results with processive mammalian DDX complexes suggest a different mechanism. We found that Lis1 has no major effect on force generation of single dynein motors bound to dynactin and does not increase the time dynein stalls on a MT before dissociating from MTs under resistive loads. Therefore, our results do not support the view that Lis1 functions primarily to regulate the tenacity of isolated dynein complexes to MTs^37, 48^. Instead, Lis1 favors recruitment of dynein to dynactin, thereby promoting the assembly of a motile complex. This finding offers an explanation for the requirement for Lis1 for transport initiation *in vivo*^38, 42^. We show that Lis1’s ability to promote the association of dynein with dynactin also favors the adoption of the two-motor state, which can account for more frequent steps and higher force generation per complex. The increased probability of recruiting a second dynein to dynactin also means that Lis1 induces more effective competition against kinesin in a tug-of-war, a result consistent with an increase in anterograde velocity observed when Lis1 is inhibited in cells^41, 43^. Remarkably, Lis1 does not have to be the part of the complex to exert its effects on motility. Lis1 dissociates from most DDX complexes before initiation of movement (Fig. 5), revealing that its primary role occurs during complex assembly.

Our results provide insights into how Lis1 enhances the affinity of dynein to dynactin. We show that this function is not dependent on reported interactions between Lis1 and the dynein tail, pointing instead to a mechanism that involves Lis1’s binding to the motor domain. Structural studies on yeast showed that Lis1 binds to the motor domain at the interface between AAA3 and AAA4 and the coiled-coil stalk^37, 47^. Assuming that mammalian Lis1 binds dynein in a similar orientation, Lis1 binding sites are positioned close to the dimerization interface on the AAA+ ring and stalk in the phi particle^26^. We propose that Lis1 binds to the open conformation of dynein and prevents switching back to the phi conformation (Fig. 6c), thereby reducing dynein autoinhibition^53^. Because the open conformation has a higher affinity to dynactin than the phi conformation^26^, Lis1 promotes the assembly of dynein with dynactin and the cargo adaptor. This mechanism would also favor the recruitment of two versus one dyneins to each dynactin.

We observed that a single dynein dimer can recruit two Lis1 dimers (Fig. 5), which is compatible with two β-propellers binding to AAA3/4 and the stalk of the same motor domain^47^. This stoichiometry might be required to prevent formation of the phi conformation. We cannot rule out the possibility that a Lis1 dimer can also form a bridge between two dyneins on dynactin in certain cases. In addition, Lis1-bound dynein may have additional structural features not present in the open conformation. Consistent with this view, we observed that mtDyn moves at the same speed as wtDyn in the absence or presence of Lis1 (Fig. 1 and Supplementary Fig. 1). High-resolution structural studies will be needed to distinguish between these possibilities. Mutagenesis studies indicated that transition between the phi-particle and open conformation is a tightly regulated process in cells^26^. Future studies are required to test whether Lis1-mediated opening of the phi conformation is also regulated by other dynein-associated proteins, such as the Lis1 binding proteins NudE and NudEL^52, 59–61^.

## Methods

### Protein expression, labeling, and purification

Human SNAPf-wtDyn, SNAPf-mtDyn, SNAPf-Dyn_LT_ (containing residues 1-1,074 of the heavy chain), BicD2N-GFP (containing residues 1-400), BicDR1-GFP, and Lis1-SNAPf were expressed in *Sf9* cells and purified using IgG affinity chromatography (using a cleavable ZZ tag), as described previously^32, 35^. *Sf9* cells were regularly tested for mycoplasma infection and no positive results were found. SNAP-tagged proteins were labeled with BG-functionalized biotin, TMR, Atto488 (NEB) or LD650 probes, and purified as described previously^35^. Dynactin was purified from pig brains using the large-scale SP-sepharose and MiniQ protocol^31^. Human Kinesin-1(1-560)-SNAPf-GFP was expressed in BL21DE3 cells and purified using Ni-NTA affinity chromatography, as described previously^62^. Concentration of isolated proteins was quantified using the Bradford colorimetric assay.

### Motility assays

Biotinylated MTs were prepared by mixing 2 μl of 5 mg/ml 2% biotin-labelled biotin with 5 μl of 10 mg/ml unlabeled pig brain tubulin in 10 μl BRB80 buffer (80 mM PIPES pH 6.8, 1mM MgCl_2_, 1 mM EGTA), followed by the addition of 10 μl polymerization buffer (2x BRB80 supplemented with 2 mM GTP and 20% anhydrous Dimethyl sulfoxide (DMSO)). Tubulin was allowed to polymerize by incubation for 40 mins at 37°C, followed by the addition of 10 nM taxol and incubation for another 40 mins. Taxol-stabilized MTs were then pelleted at 20,000 g for 12 min and resuspended in BRB80 buffer containing 10 nM taxol and 1 mM Dithiothreitol (DTT).

Motility chambers were made by applying two strips of double-sided tape on a glass slide, and then placing a clean coverslip on the top. The glass surface was first passivated by BSA and functionalized with biotin by flowing 20 μl of 1 mg/ml BSA-biotin (Sigma), followed by washing with 40 μl of dynein motility buffer (DMB: 30 mM HEPES pH 7.0, 5 mM MgSO_4_, 1 mM EGTA, 1 mM TCEP (tris(2-carboxyethyl)phosphine) supplemented with1.25 mg/ml casein (Sigma). To immobilize biotinylated MTs on the functionalized surface, 20 μl of 1 mg/ml streptavidin (NEB) was then flowed through and washed with 40 μl of the same buffer.

For DDB, DDR, and DTR motility, 0.7 μl of 1.2 mg/ml LD650-labeled dynein was mixed with 1 μl of 1.6 mg/ml dynactin, and 1.5 μl of 3-4 mg/ml cargo adaptor (BicD2N-GFP or BicDR1-GFP) in 10 μl DMB supplemented with 1 mg/ml Bovine Serum Albumin (BSA). For DTR experiments, 1 μl of 3.7 mg/ml TMR-labeled dynein tail was added to the mixture. For co-localization experiments, 0.9 μl of 1 mg/ml TMR-labeled dynein was additionally included in the motility mix. For single-color Lis1 experiments, 1 μl of 0.9 mg/ml TMR-labeled Lis1 was added to 0.7 μl of 1.2 mg/ml LD650-labeled dynein. For two-color Lis1 experiments, 0.5 μl of 0.9 mg/ml TMR-labeled Lis1 and 0.7 μl of 0.9 mg/ml Cy5-labeled Lis1 were added to 1.2 μl of 0.8 μl of unlabeled dynein and 0.5 μl of 3.3 mg/ml Atto488-labeled Dyn_LT_. The complexes were incubated on ice for 10 mins, diluted in DMB supplemented with 1.25 mg/ml casein (DMB-C), and flowed into the chamber. The motility mix was kept for 2 mins and then washed with 40 μl of DMB-C. To record motility, 20 μl of dynein stepping buffer (DMB-C supplemented with 1 mM Mg.ATP, 2.5 mM PCA (protocatechuic acid), 35 μg/ml PCD (protocatechuate-3,4-dioxygenase)) was introduced into the chamber and the sample immediately imaged for 3 mins at 23°C. For experiments with Lis1, 1 μl of 1.3 mg/ml Lis1-SNAPf was included in the reaction mixture and an additional 1 μl of the same protein was then reintroduced into the motility mix with the dynein stepping buffer.

Single-molecule motility experiments were performed using a custom-built TIRF microscope equipped with a 100x 1.49 N.A. apochromat oil-immersion objective (Nikon) and perfect focusing system on an inverted microscopy body (Nikon Ti-Eclipse). Fluorescence signal was detected using an electron-multiplied charge-coupled device (EM-CCD) camera (Andor, Ixon EM^+^). The sample that contained LD650 was excited with 0.05 kW cm^−2^ 633 laser beam (Coherent), and emission signal was filtered using 655/40 nm bandpass emission filter (Semrock). Movies were recorded using an effective pixel size of 160 nm at 300 ms per frame. For two- and three-color fluorescence assays, imaging was performed on a multi-color TIRF microscope (Nikon) using alternating excitation and time-sharing mode of emission collection. Atto488-, TMR- and LD650-labeled samples were excited using 0.05 kW cm^−2^ 488, 532 and 633 nm laser beams (Coherent) and fluorescence signals was detected on an ImagEM X2 EM-CCD camera (Hamamatsu). Movies were recorded at 150 ms per frame per color for two-color and at 100 ms per frame per color for three-color fluorescence assays. Effective pixel size was 108 nm.

Kymographs were generated from movies in ImageJ. Processive movement and velocity were then defined and measured, as described previously^51^. For two color imaging, the two channels were overlaid using the merge function. Resulting kymographs were then manually scored for processive events that show co-localization between the two channels. Labeling efficiency of a dynein dimer with at least one TMR or LD650 was 96%, as determined by spectrophotometry. The fractions of the complexes containing two dyneins was calculated using the TMR-Dyn and LD650-Dyn colocalization measurements, after accounting for unlabeled complexes, and complexes assembled with two dyneins labeled with the same color^35^.

### High resolution fluorescence-tracking assays

QDs were functionalized with benzyl guanine by mixing 5 μl of 8 μM amino (PEG) QDs emitting at 655 nm (ThermoFisher) with 2 μl of 20 mM BG-GLA-NHS (NEB) in 20 μl 100 mM sodium borate buffer, pH 8.0 for 40 min at room temperature. To remove excess BG-GLA-NHS, functionalized QDs were concentrated through five consecutive spins through 100,000 MWCO centrifugal filter units (Amicon). Finally, spin-concentrated QDs were suspended in 100 μl DMB and stored at 4°C.

For tracking the motility of individual dynein complexes, 0.5 μl of 1.7 mg/ml SNAPf-dynein was mixed with 1 μl of 1.6 mg/ml dynactin, and 1.5 μl of 3.5 mg/ml BicD2N-GFP, and 1 μl of 1.3 mg/ml Lis1-SNAPf in 10 μl DMB supplemented with 1 mg/ml BSA. The complex was incubated in ice for 10 mins followed by the addition of 1 μl of 400 nM BG-functionalized QDs for another 10 mins in ice. The mixture was then diluted in DMB-C and flowed into the motility chamber for 2 mins, followed by washing with 80 μl of DMB-C. 20 μl of dynein stepping buffer containing 2 μM Mg.ATP was flowed into the chamber and motility was immediately recorded. For tracking the stepping of dynein in the presence of Lis1, 1 μl of 1.3 mg/ml Lis1-SNAPf was included in the dynein stepping buffer. The sample was excited with a 1 kW cm^−2^ 488 nm beam (Coherent) and movies were recorded at 30 ms per frame on Ixon EM+ EM-CCD camera (Andor). For stepping analysis, fluorescence spots of QDs were localized using a 2-dimenstional Gaussian fitting and the resulting trajectories fitted into steps using a custom-written algorithm based on Schwartz Information Criterion ^63, 64^.

### Tug-of-war assays

To prepare a DNA tether between DDB and kinesin, two complementary DNA strands were first functionalized with benzyl guanine as described previously. Briefly, 10 μl of 100 μM DNA oligos containing an amino group modification at their 5’ends were mixed with 8 μl of 20 mM BG-GLA-NHS (NEB) in 40 μl of 50 mM HEPES buffer containing 50% anhydrous DMSO, pH 8.5. The reaction was kept overnight at room temperature. Excess unreacted ligand was removed and the BG-functionalized oligos was purified by ethanol precipitation of DNA. Finally, isolated DNA was dissolved in 50 μl DMB and stored at 4°C. Concentration of BG-DNA was estimated from the absorbance at 260 nm.

BicD2N-SNAPf and kinesin-SNAPf-GFP were labeled with BG-functionalized oligos by mixing protein with DNA in DMB for 1 hr at 4°C. DNA and protein concentrations were optimized to yield ~ 30% efficiency of protein labeling to ensure that the likelihood of dual labeling of a single dimeric protein with two DNA oligos was minimized (<9%). The labeling efficiency was quantified by comparing the intensities of labeled to unlabeled bands on 4-12% Bis-Tris SDS-PAGE (Invitrogen). Excess unreacted DNA was removed from BicD2N-SNAPf using a TSKgel G4000SWXL size exclusion column (Tosoh). In case of kinesin, 10-fold molar excess of BG-GLA-TMR (NEB) was added to the motor-DNA mixture and the sample was incubated for additional 30 min at 4°C. Excess DNA and dye were removed by a MT bind and release assay.

In Tug-of-war experiments, 1 μl of 1.2 mg/ml DNA-labeled BicD2N-SNAPf was mixed with 0.7 μl of 1.2 mg/ml LD650-labeled dynein and 1 μl of 1.6 mg/ml dynactin in 10 μl DMB supplemented with 1 mg/ml BSA. For experiments with Lis1, 1 μl of 1.3 mg/ml Lis1-SNAPf was added to the mixture. The mixture was incubated in ice for 10 mins, followed by the addition of 1 μl of 0.5 M NaCl and 0.5 μl of 0.8 mg/ml DNA- and TMR- labeled kinesin-SNAPf-GFP, and incubation on ice for a further 20 mins. Proteins were then diluted in DMB-C and flowed into the chamber, followed by washing with 80 μl of DMB-C and imaging in 20 μl dynein stepping buffer supplemented with 1 mM ATP and 1 μl of 1.3 mg/ml Lis1-SNAPf in case of Lis1 experiments.

### Optical trapping assays

For DDB and DDR experiments, complexes assembled with SNAPf-dynein, dynactin, and BicD2N-GFP or BicDR1-GFP were mixed with 860 nm diameter anti-GFP coated latex beads in ice for 10 mins. Carboxy latex beads were functionalized with anti-GFP antibodies, as described previously^65^. Briefly, 200 μl of 860 nm latex beads (Life Technologies) were washed and resuspended in activation buffer (10 mM MES, 100 mM NaCl, pH 6.0) and then coated by mixing with ~ 2 mg of custom-made polyclonal rabbit anti-GFP antibodies (Covance) in 100 μl activation buffer supplemented with 1 mg each of N-hydroxysulfosuccinimide (Sulfo-NHS) and 1-Ethyl-3-(3-dimethylaminopropyl) carbodiimide (EDC) crosslinkers (Pierce) dissolved in 100 μl of dimethylformamide (DMF). Finally, the beads were passivated with BSA, washed and stored in 200 μl phosphate-buffered saline (PBS) supplemented with 0.5 mg/ml BSA and 0.1% sodium azide at 4 °C. For DTR experiments, SNAPf-Dyn_LT_ was labeled with biotin, and biotin-Dyn_LT_, mtDyn, dynactin and BicDR1 were incubated with 800 nm diameter streptavidin-coated beads (Spherotech) in ice for 10 mins. The protein-coated beads were then diluted in DMB-C and flowed into the motility chamber in dynein stepping buffer supplemented with 1 mM ATP. The protein concentration in the mixture was gradually reduced in assays until less than 30% of the tested beads exhibited motility activity in contact with Cy5-labeled axonemes to ensure that >95% of the beads are driven by a single complex. For force measurements in the presence of Lis1, 600 nM Lis1-SNAPf was added the bead-protein mixture and also later added to the stepping buffer.

Force measurements were performed on a custom-built optical trap on a Nikon Ti-Eclipse microscope body consisting of a 2 W 1,064 nm continuous wave laser beam (Coherent) and a 100 × 1.49 NA Plan-Apo objective (Nikon), as described previously ^65^. Beads were trapped by the laser beam steered by two computer-controlled perpendicular acousto-optical deflectors (AA Electronics). The sample was excited with a 633 nm laser (Coherent) and Cy5-labeled axonemes were imaged using a monochrome camera (The Imaging Source). To detect bead position relative to the center of the trap, a position-sensitive detector (First Sensor Inc.) was placed at the back focal plane of a 1.4 N.A. oil-immersion condenser (Nikon). Trap stiffness was derived by fitting the power spectrum of a trapped bead that was rapidly raster scanned in both x and y directions using the acousto-optical deflectors to a Lorentzian spectrum. The typical spring constant used in these experiments was ~ 0.04 pN/nm to allow motors to travel 100-150 nm before stalling. The PSD data were recorded at 20 kHz during calibration and the resulting curve was fit to a cubic polynomial to calibrate the response of the PSD in each sample. For fixed trap assays, PSD data were collected at 5 kHz and downsampled to 500 Hz for ease of visualization. To qualify as a stall event, the bead position should remain stationary for at least 100 ms before rapid (<2 ms) jumping towards the trap center, implicating release of the motor from the MT. Stall force histograms are then generated from individual stall events that were manually scored. For force-clamps assays, PSD signal was acquired at 5 kHz and position feedback was performed at 100 Hz. Beads that walked for at least 100 nm were subjected to force feedback and resulting runs were downsampled to 500 Hz and fit to a step-finding algorithm as described previously^65^. Force-clamp runs that are shorter than 200 nm or included instant jumps larger than 50 nm were excluded from the analysis.

### Statistics and reproducibility

At least three independent experiments were performed to obtain each result. The exact number of replicates (n) of every dataset is given at the corresponding figure legends. Statistical analysis methods are stated in the main text or the figure legend.

## Code availability

Codes used in this paper are available from the corresponding author upon request.

## Data availability

All data that support the conclusions are available from the authors on request.

## Authors contribution

M.M.E., J.B., S.L.B. and A.Y. conceived the study and designed the experiments. M.M.E. purified dynein, dynactin and cargo adaptors. J.B. purified Lis1 proteins. M.M.E. and E.K. labeled the proteins with DNA and fluorescent dyes and performed the single molecule motility experiments. M.M.E. and E.K. performed fluorescent tracking assays. M.M.E., S.V. and E.K. performed optical trapping assays. M.M.E., S.L.B. and A.Y. wrote the manuscript, and all authors read and edited the manuscript.

## Acknowledgments

We are grateful to the members of the Yildiz laboratory for helpful discussions. This work was funded by grants from the NIH (GM094522), and NSF (MCB-1055017, MCB-1617028) to A.Y., the MRC (MC_U105178790) to S.L.B., and a DFG research fellowship (BA5802/1-1) to J.B.

