## Supplementary Information for "Lis1 activates dynein motility by pairing it with dynactin"

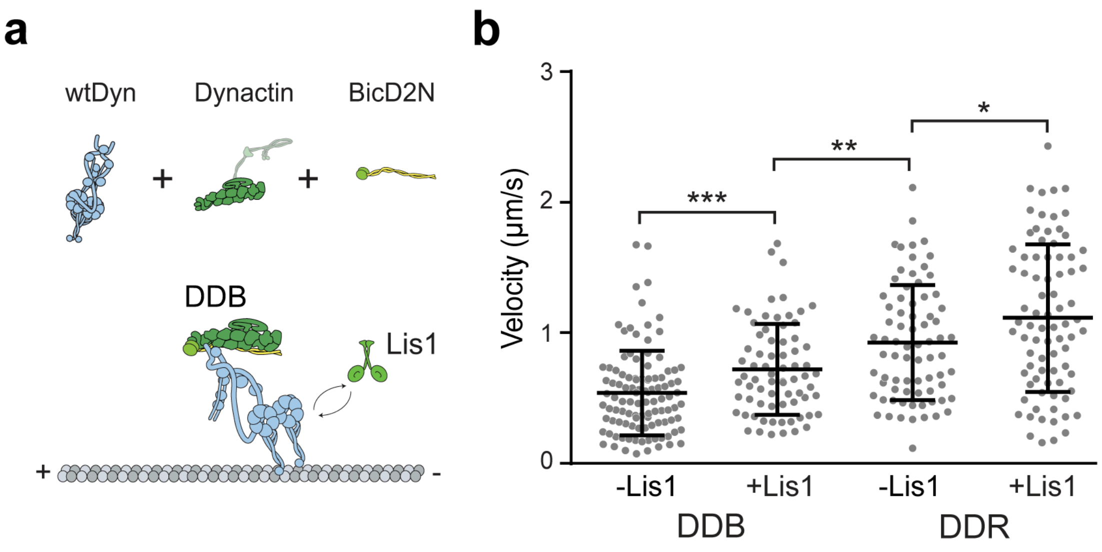


**Supplementary Figure 1. Lis1 increases the velocity of complexes assembled with wild-type dynein.** **(a)** Assembly of DDB and DDR with wtDyn. **(b)** Velocity distribution of DDB and DDR complexes assembled in the presence and absence of 600 nM Lis1. The line and whiskers represent the mean and s.d., respectively ($n$ = 106, 72, 75, and 81 from left to right, ***p < 0.005, **p <0.001, *p <0.05, two-tailed t-test).

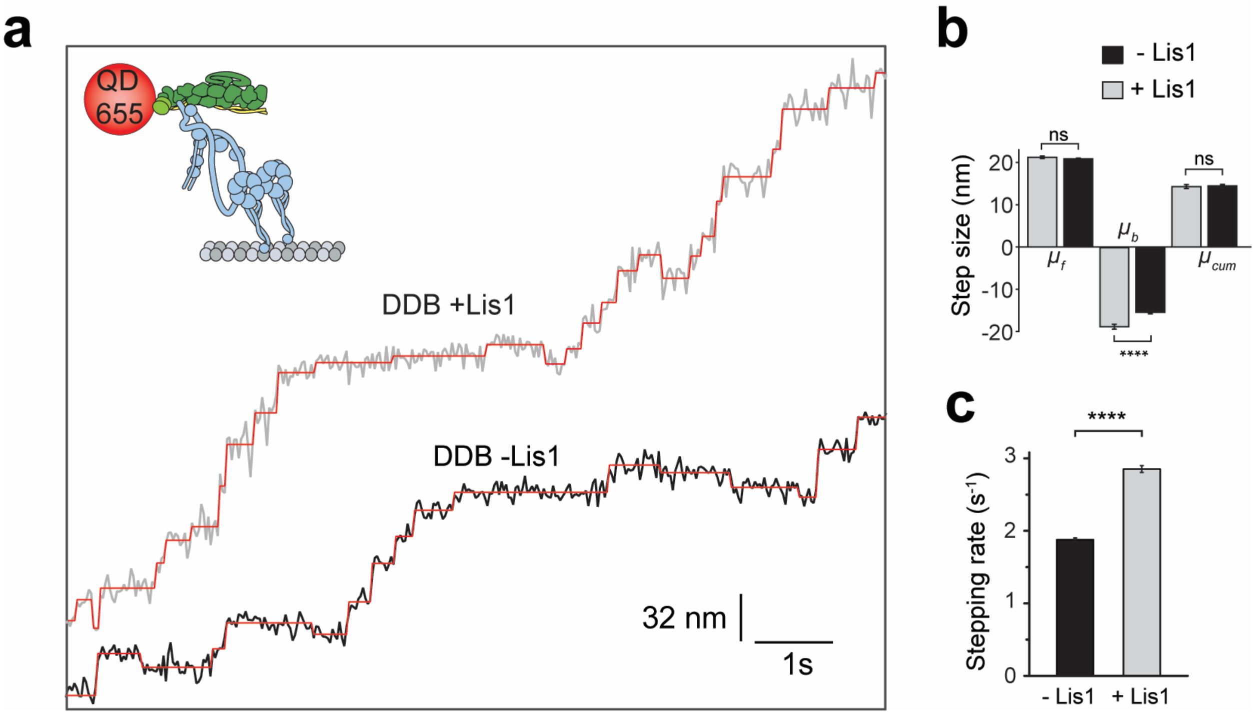


**Supplementary Figure 2. Step analysis of DDB in the presence and absence of Lis1. (a)** Additional examples of DDB stepping in the presence and absence of 600 nM Lis1. **(b)** The average size of steps taken in forward ($\mu_f$), backward, ($\mu_b$), and both ($\mu_{cum}$) directions along the longitudinal axis of the MT (ns: non-significant, ****$p < 0.0001$, two-tailed t-test). Error bars are s.e.m. **(c)** Stepping rates estimated from the exponential fit in Figure 1f (****$p < 0.0001$, two-tailed t-test). Error bars are s.e of the fit.

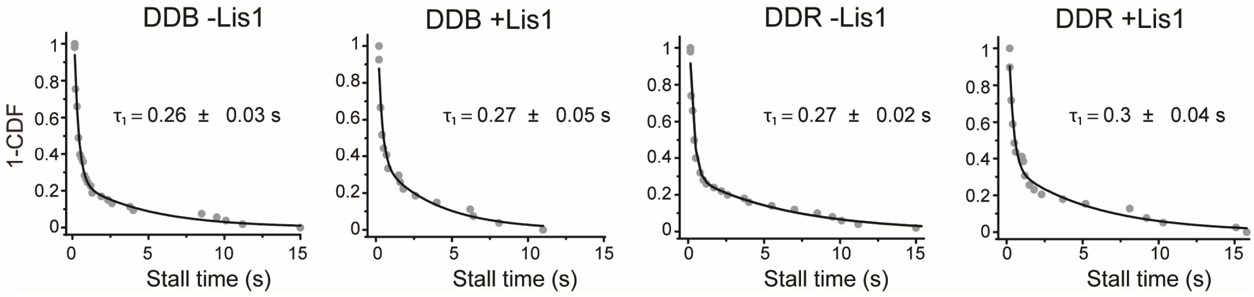


**Supplementary Figure 3. Lis1 does not increase the stall duration of dynein bound to dynactin and a cargo adaptor.** Stall time distribution of DDB and DDR in absence and presence of Lis1. Inverse cumulative distribution of stall durations. Solid curves represent fitting to a two-exponential decay (decay time ± s.e., *n* = 53, 27, 50, and 39 from left to right).

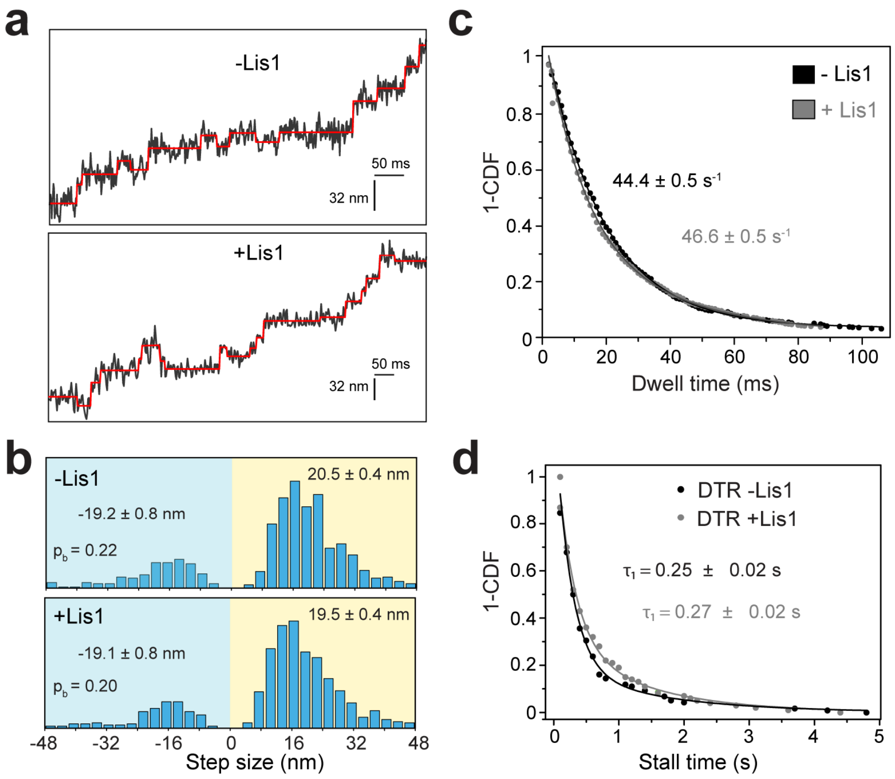


**Supplementary Figure 4. Lis1 does not affect stall time and stepping behavior of single dynein. (a)** Streptavidin beads are sparsely coated with **biotin-**$Dyn_{LT}$ in the presence of mtDyn, dynactin and BicDR1, and trapped with a focused laser beam. Example traces of beads driven by DTR in the presence and absence of Lis1 against 1 pN hindering force. The raw stepping data are shown in black and steps fitting are in red. **(b)** Normalized histograms of DTR steps taken in the longitudinal direction (*n* = 729 for -Lis1 and 718 for +Lis1). Average sizes of steps taken in forward and backward directions (± s.e.m.) and the probability of backward stepping in the presence and absence of Lis1 are statistically indistinguishable (p = 0.6, two-tailed t-test). **(c)** Distribution of dwell times between consecutive steps along the longitudinal axis of the MT. A fit to an exponential decay reveals the decay rate (rate ±s.e., *n* = 734 for DTR -Lis1 and 725 for DTR +Lis1). **(d)** Inverse cumulative distribution of stall durations of DTR in the presence and absence of Lis1. Solid curves represent fitting to a two-exponential decay (decay time ± s.e., n = 118 for DTR - Lis1 and 100 for DTR + Lis1).

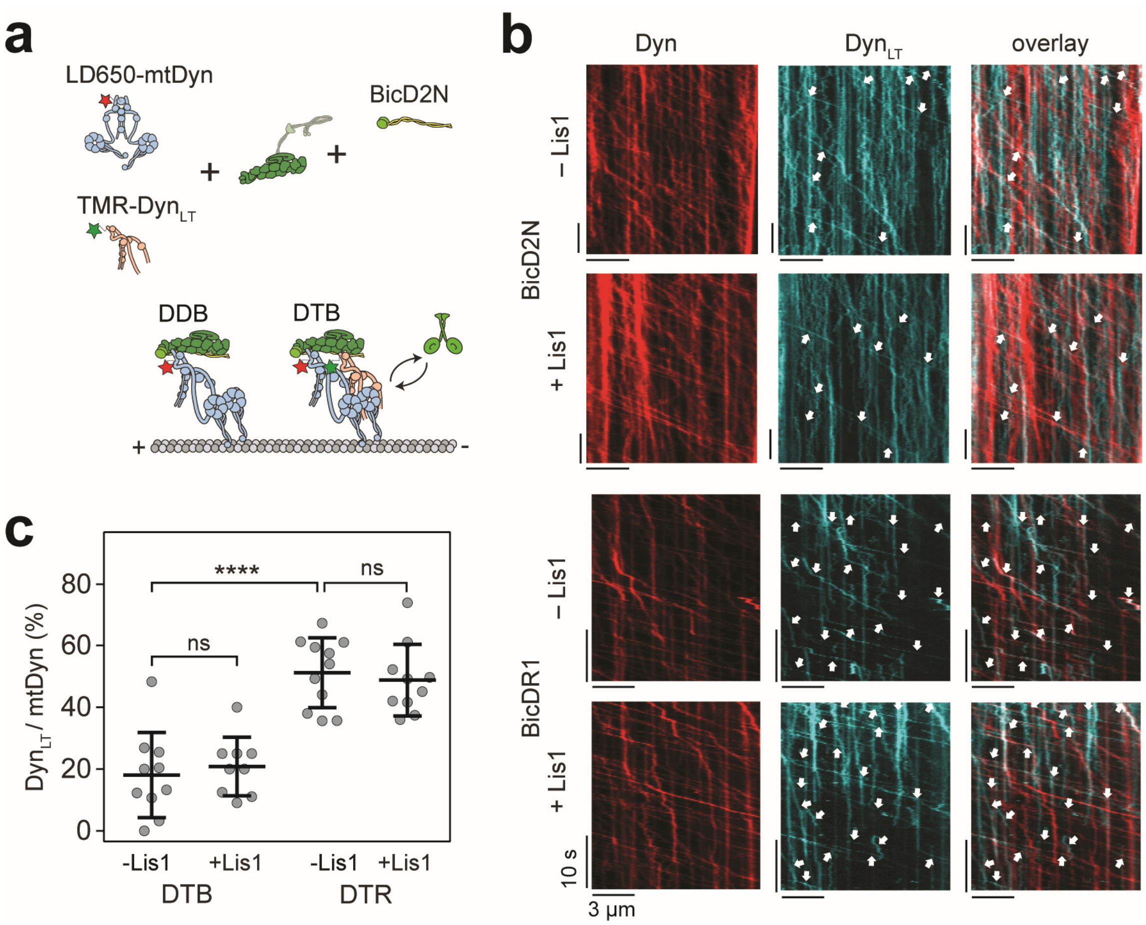


**Supplementary Figure 5. Lis1 does not increase recruitment of dynein tail to dynactin. (a)** Schematic shows the assembly of DDB and DTB complexes using TMR-Dyn$_{LT}$, LD650-mtDyn, dynactin and BicD2N. **(b)** Representative kymographs show the motility of LD650-Dyn and TMR-Dyn$_{LT}$ assembled with BicD2N or BicDR1 in the presence and absence of Lis1. White arrows point to complexes that contain both LD650-mtDyn and TMR-Dyn$_{LT}$. **(c)** The ratio of processive runs by TMR-Dyn$_{LT}$ to LD650-mtDyn on individual MTs in the presence and absence of Lis1. The line and whiskers represent the mean and s.d., respectively ($n$ = 10, 9, 11, and 10 MTs from left to right, ns: non-significant, ****p <0.0001, two-tailed t-test).

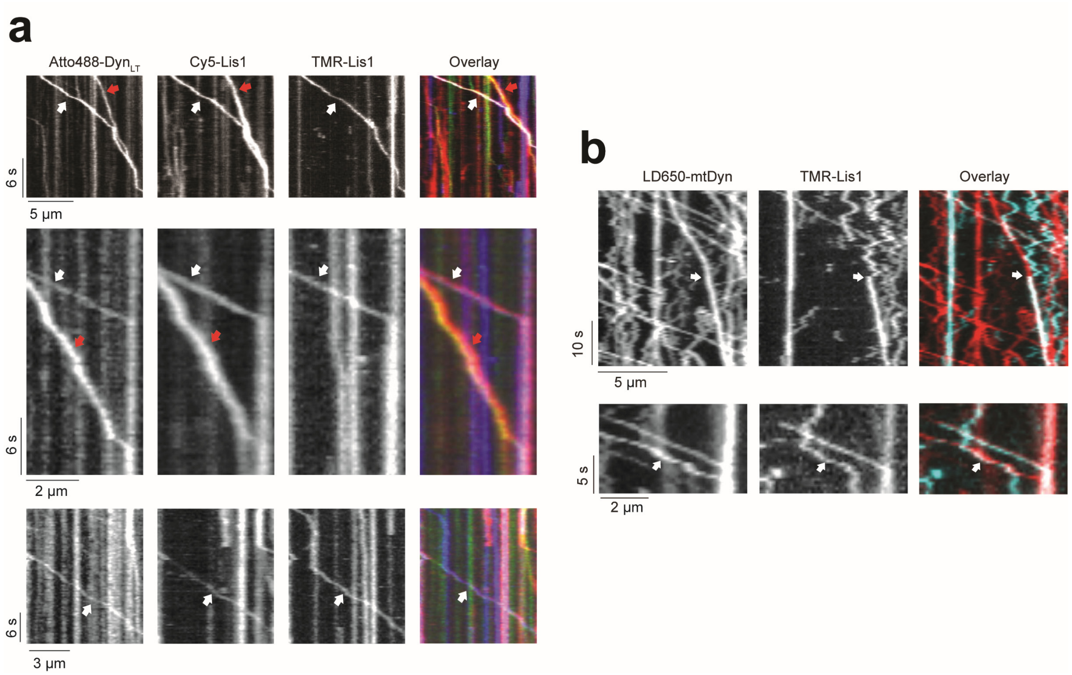


**Supplementary Figure 6. Additional examples of binding events of Lis1 to DDX during processive movement. (a)** Additional kymographs show single- and dual Lis1 binding to motile DTR complexes. Red arrows represent colocalization of Atto488-Dyn$_{LT}$ (green) and Cy5-Lis1 (red). White arrows represent colocalization of Atto488-Dyn$_{LT}$ (green) with both Cy5-Lis1 (red), and TMR-Lis1 (cyan). **(b)** Rare events of dynamic binding of Lis1 to dynein as DDB walks along an MT. White arrows represent colocalization of LD650-Dyn (red) and TMR-Lis1 (cyan). In the top kymograph, Lis1 initially diffuses on an MT and then binds to DDB during processive movement. Lis1 binding reduces the velocity of the complex. In the bottom kymograph, a diffusing Lis1 initially binds and later dissociates from DDB, without significantly affecting the velocity of the complex.

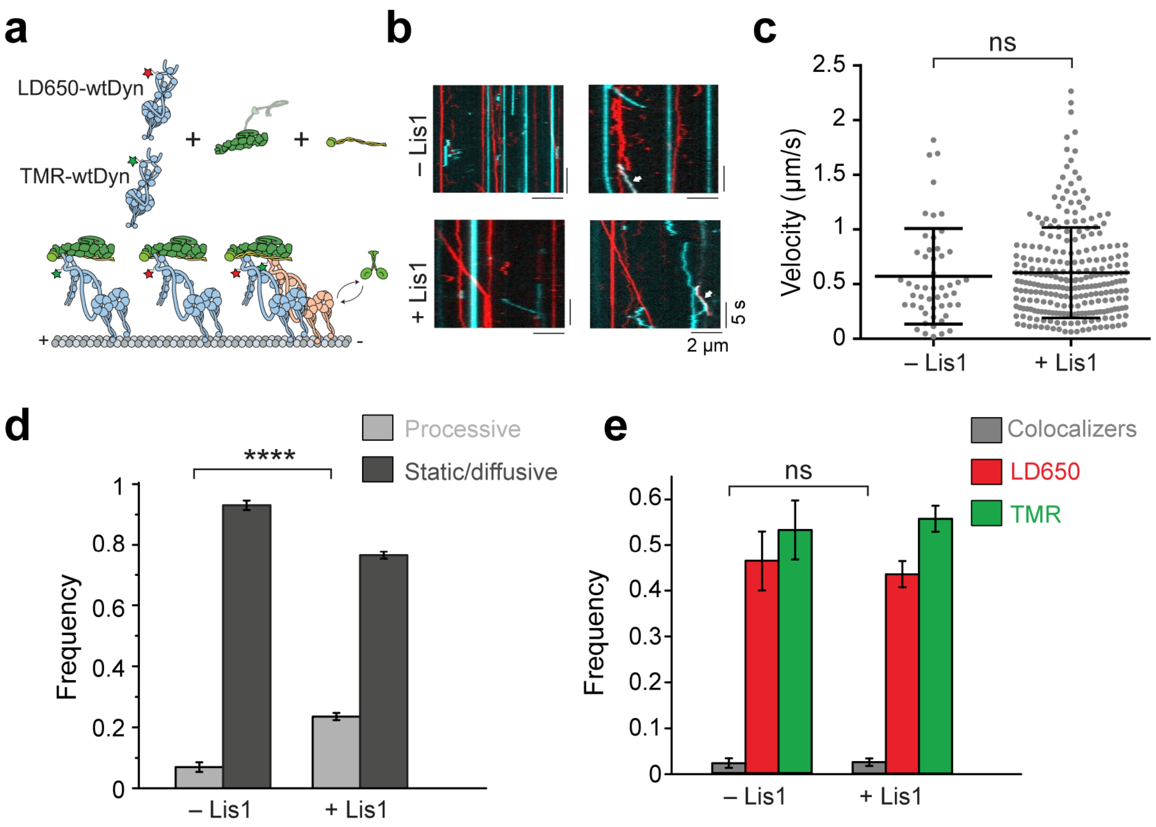


**Supplementary Figure 7. At limiting dynein concentration, Lis1 recruits a single dynein to dynactin and BicD2N. (a)** Schematic depiction of DDB assembly using 5 nM LD650-wtDyn and TMR-wtDyn in the absence and presence of Lis1. **(b)** Representative kymographs show the motility of LD650- (red) and TMR- (cyan) wtDyn. Left kymographs show single-colored runs and right kymographs show rare events of TMR-LD650 colocalization (white arrows). **(c)** Velocity distribution of DDB motility with and without 600 nM Lis1. The center line and whiskers represent the mean and s.d., respectively ($n$ = 51 and 257 from left to right, ns: non-significant, two-tailed t-test). **(d)** Fraction of processive and static/diffusive DDB complexes on MTs (mean ± s.e.m, $n$ = 59, 788, 303 and 984 from left to right, ****p < 0.0001, two-tailed t-test). **(e)** Fraction of processive complexes that contain TMR, LD650, and TMR-LD650 colocalizers (mean ± s.e.m, $n$ = 59 for -Lis1 and 303 for + Lis1, ns: non-significant, two-tailed t-test).

**Supplementary Movie Legends:**

**Supplementary Movie 1. Motility of single DDBs along MTs at 1 mM ATP.** LD650-mtDyn was assembled with dynactin and BicD2N and motility along surface-attached MTs in the absence and presence of Lis1 was imaged under TIRF illumination. Scale bar is 5 μm. Stop watch shows time in seconds.

**Supplementary Movie 2. Motility of single DDRs along MTs at 1 mM ATP.** LD650-mtDyn was assembled with dynactin and BicDR1 and motility along surface-attached MTs in the absence and presence of Lis1 was imaged under TIRF illumination. Scale bar is 5 μm. Stop watch shows time in seconds.

**Supplementary Movie 3. Motility of DDB-kinesin co-localizers.** LD650-mtDyn was assembled with dynactin and BicD2N. LD650-DDB (red) and TMR-kinesin (cyan) were tethered using a DNA scaffold and motility along surface-attached MTs in the absence and presence of 600 nM Lis1 was imaged with two-color TIRF illumination. Colocalizers are denoted by white arrows. Scale bar is 5 μm. Stop watch shows time in seconds.

**Supplementary Movie 4. Motility of single DTRs along MTs at 1 mM ATP.** LD650-mtDyn (red) and TMR-$Dyn_{LT}$ (cyan) were mixed in the presence of dynactin and BicDR1. Motility along surface-attached MTs in the absence and presence of 600 nM Lis1 was imaged with two-color TIRF illumination. Colocalizers that move along a single MT are denoted by white arrows. Scale bar is 5 μm. Stop watch shows time in seconds.

**Supplementary Movie 5. Recruitment of two dyneins to dynactin by BicD2N and BicDR1.** LD650-mtDyn (red) and TMR-mtDyn (cyan) were mixed with dynactin and a cargo adaptor in the absence and presence of 600 nM Lis1. Motility of DDB and DDR complexes along surface-attached MTs were imaged with two-color TIRF illumination. Colocalizers along a single MT are denoted by white arrows. Scale bar is 5 μm. Stop watch shows time in seconds.

**Supplementary Movie 6. Colocalization of two Lis1 dimers to DTR.** Atto488-$Dyn_{LT}$ (green), TMR-Lis1 (blue), and LD650-Lis1 (red) were mixed with dynactin and BicDR1. Motility along surface-attached MTs was imaged with three-color TIRF illumination. Atto488, TMR, LD650, and overlaid frames are separately shown for ease of visualization. Colocalizers are denoted by white arrows. Scale bar is 3 μm. Stop watch shows time in seconds.

**Supplementary Movie 7. Motility of single DDBs along MTs at limiting dynein concentration.** 5 nM of LD650-mtDyn or LD650-wtDyn was mixed with dynactin and BicD2N in the absence and presence of 600 nM Lis1. Motility along surface-attached MTs was imaged under TIRF illumination. Scale bar is 5 μm. Stop watch shows time in seconds.
